# Crosstalk in oxygen homeostasis networks: SKN-1/NRF inhibits the HIF-1 hypoxia-inducible factor in *Caenorhabditis elegans*

**DOI:** 10.1101/2021.03.12.435071

**Authors:** Dingxia Feng, Zhiwei Zhai, Zhiyong Shao, Yi Zhang, Jo Anne Powell-Coffman

**Affiliations:** Department of Genetics, Development and Cell Biology, Iowa State University, Ames, Iowa, United States of America

## Abstract

During development, homeostasis, and disease, organisms must balance responses that allow adaptation to low oxygen (hypoxia) with those that protect cells from oxidative stress. The evolutionarily conserved hypoxia-inducible factors are central to these processes, as they orchestrate transcriptional responses to oxygen deprivation. Here, we employ genetic strategies in *C. elegans* to identify stress-responsive genes and pathways that modulate the HIF-1 hypoxia-inducible factor and facilitate oxygen homeostasis. Through a genome-wide RNAi screen, we show that RNAi-mediated mitochondrial or proteasomal dysfunction increases the expression of hypoxia-responsive reporter *Pnhr-57:GFP* in *C. elegans.* Interestingly, only a subset of these effects requires *hif-1.* Of particular importance, we found that *skn-1* RNAi increases the expression of hypoxia-responsive reporter *Pnhr-57:GFP* and elevates HIF-1 protein levels. The SKN-1/NRF transcription factor has been shown to promote oxidative stress resistance. We present evidence that the crosstalk between HIF-1 and SKN-1 is mediated by EGL-9, the prolyl hydroxylase that targets HIF-1 for oxygen-dependent degradation. Treatment that induces SKN-1, such as heat, increases expression of a *Pegl-9:GFP* reporter, and this effect requires *skn-1* function and a putative SKN-1 binding site in *egl-9* regulatory sequences. Collectively, these data support a model in which SKN-1 promotes *egl-9* transcription, thereby inhibiting HIF-1. We propose that this interaction enables animals to adapt quickly to changes in cellular oxygenation and to better survive accompanying oxidative stress.

## Introduction

Oxygen homeostasis has profound effects on health and fitness. Oxygen serves as the terminal electron acceptor in the oxidative phosphorylation processes that generate energy for life. When oxygen levels are low (hypoxia), cells and tissues must adapt quickly by increasing oxygen delivery, adjusting the levels of key metabolic enzymes, and limiting the accumulation of misfolded proteins. While oxygen is essential, it is also highly reactive. The reactive oxygen species (ROS) generated by cellular metabolism and signaling processes can damage macromolecules, and excess ROS are thought to contribute to cellular aging and deterioration [1–3]. One of the central challenges of aerobic life is to coordinate the biological networks that control disparate aspects of oxygen homeostasis.

This balance between surviving hypoxic stress and mitigating the potential damage caused by reactive oxygen species is especially important in cardiovascular development and disease. When ischemia blocks circulation to a mammalian tissue, oxygen levels drop, and cells induce hypoxia-inducible transcription factors (HIFs). Upon reperfusion and reoxygenation of the tissue, mammalian cells respond by rapidly degrading HIF and inducing the NRF2 transcription factor [4, 5]. However, intermittent hypoxia has been shown to induce both HIF-1α and NRF2 [6, 7]. NRF2 activates phase II detoxification genes to mitigate the effects of oxidative insults [8, 9]. Although mammalian HIF and NRF2 share some common target genes such as aldehyde dehydrogenase 1A1 or heme oxygenase-1 HO-1, the genes induced by re-oxygenation are largely distinct from those that respond to oxygen deprivation [5, 10, 11]. Crosstalk between these two pathways is complex and context specific in mammals [12], as these important transcription factors facilitate the rapid changes in gene expression needed to limit reperfusion injury and regulate oxygen-dependent developmental processes.

*C. elegans* has been proven to be an excellent model system for studying the regulatory networks that govern oxygen homeostasis. The *C. elegans* genome encodes a single hypoxia-inducible factor alpha subunit (HIF-1), and the *hif-1* gene has been shown to have important roles in stress response and in aging [13–17]. HIF protein levels and HIF activity are tightly regulated. When oxygen is abundant, the HIF alpha subunit is hydroxylated by the PHD/EGL-9 enzymes. Once modified, HIFα protein interacts with the Von Hippel-Lindau tumor suppressor (VHL) and is targeted for ubiquitination and proteasomal degradation [18–23]. Thus, in hypoxic conditions, HIF-1 protein is stable, and the transcription factor complex can activate the expression of a battery of genes that enable adaptation to low oxygen. This pathway for oxygen-dependent degradation of HIF protein is evolutionarily conserved. The *C. elegans hif-1, aha-1, egl-9,* and *vhl-1* genes are orthologous to mammalian *HIFα, HIFβ, PHD,* and *VHL,* respectively [24–26]. The targets of *C. elegans* HIF-1 include *egl-9* and *rhy-1,* genes that inhibit HIF-1 expression and activity [27–30]. In wild-type animals, these negative feedback loops keep HIF-1 activity in check and limit the potentially adverse effects of HIF-1 over-activation.

The *C. elegans skn-1* gene is homologous to mammalian *NRF1/2/3* [31]. SKN-1 regulates the expression of a battery of genes with cytoprotective functions, including phase II detoxification genes [32, 33]. SKN-1 is activated by a range of stresses or toxicants that cause oxidative stress, and SKN-1 promotes resistance to these insults [32, 34–37].

Here, we investigate the cellular processes and transcriptional networks that regulate HIF-1 function. We describe an unbiased genetic screen to identify genes that inhibit *C. elegans* HIF-1. We discover that SKN-1/NRF represses HIF-1 protein levels. Hence, SKN-1-mediated repression of HIF-1 may provide a mechanism by which cells can rapidly respond to specific environmental stresses and optimize gene expression to achieve oxygen homeostasis. We investigate the hypothesis that this cross talk is mediated by EGL-9, the oxygen-sensing prolyl hydroxylase that modulates HIF-1 stability and activity.

## Results

To identify genes and cellular processes that attenuated HIF-1-mediated gene expression, we conducted a genome-wide RNAi screen. This experimental strategy relied on the *Pnhr-57:GFP* reporter gene, which had been shown to be responsive to HIF-1 and hypoxia [28, 29]. Through chromatin immunoprecipitation experiments, we confirmed that *nhr-57* was a direct target of HIF-1 (S1 Fig). We screened a bacterial RNAi library representing ~80% of *C. elegans* genes [38], and we identified 179 genes for which RNAi increased *Pnhr-57:GFP* expression, as assayed by inspection under a fluorescent stereomicroscope (screen design illustrated in Fig 1A). These genes are listed in S1 Table and their functions are summarized in Fig 1B. Remarkably, 89 genes were predicted to have mitochondrial or metabolic functions, and the majority of these were electron transport chain components (42 genes) or subunits of mitochondrial ribosomes (24 genes). The second largest category of the genes was protein folding or turnover (33 genes), and the majority of these were proteasomal components. The remaining major categories include vesicular transport, transporters and channels, signaling and cytoskeleton, transcription and DNA or RNA processing.

**Fig 1.**
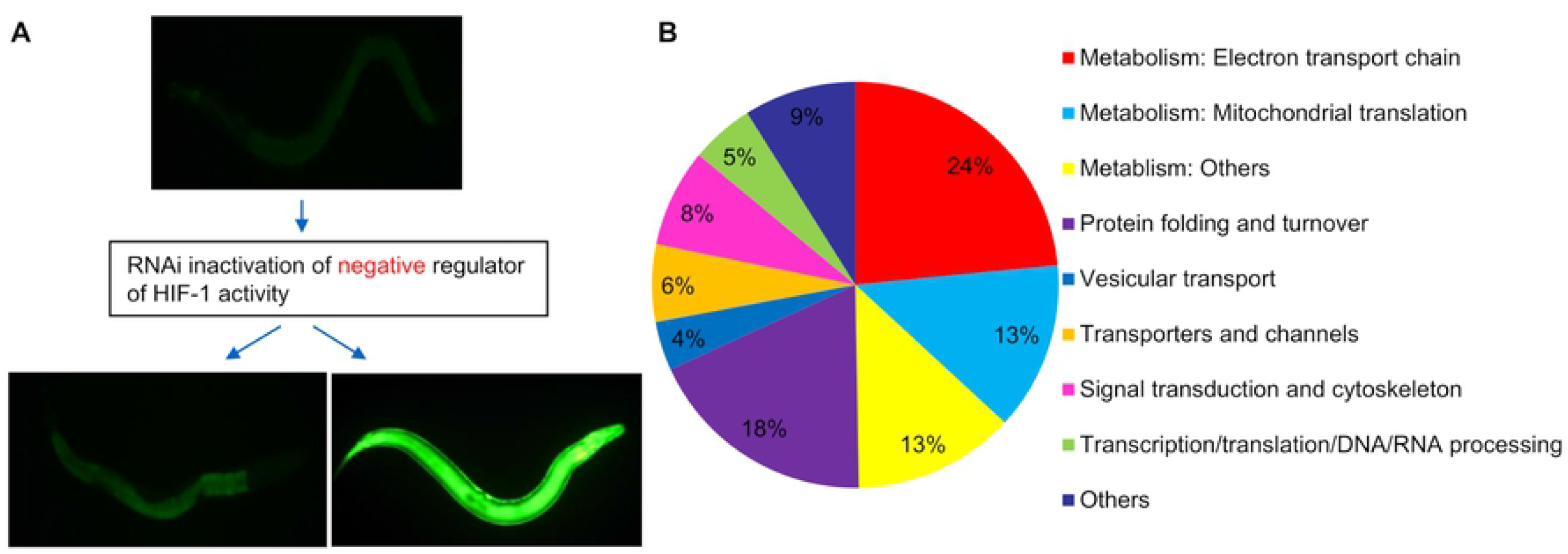
Genome-wide RNAi screen to identify negative regulators of HIF-1-mediated gene expression. (A) Illustration of screen design. The screen identified RNAi treatments that increased the expression of *Pnhr-57:GFP,* a reporter that has been shown to be induced by hypoxia and positively regulated by HIF-1. (B) Functional summary of the 179 genes identified from the RNAi screen.

Recognizing that most eukaryotic genes are coordinately regulated by multiple transcription factors, we did a secondary screen to identify those RNAi treatments that had a clear *hif-1*-dependent effect on *Pnhr-57:GFP* expression. To do this, we compared the *Pnhr-57:GFP* induction of these 179 RNAi treatments in wild-type animals and *hif-1*-deficient animals. Most of the 179 RNAi treatments increased the expression of *Pnhr-57:GFP* independent of *hif-1.* However, the *Pnhr-57:GFP* induction by 13 RNAi treatments showed a strong *hif-1*-dependent effect (S2 Table). Among these 13 genes, as expected, RNAi for *egl-9, rhy-1,* and *vhl-1,* previously characterized negative regulators of *C. elegans* HIF-1 [26, 29], increased *Pnhr-57:GFP* expression in wild-type animals, but not in *hif-1* mutants. These results validated the efficacy of our screen approach, and gave us the confidence to continue investigating the potentially new negative regulators of HIF-1 among these 13 genes.

### *skn-1* attenuates HIF-1 protein levels and HIF-1 function

We were especially intrigued by the finding that *skn-1* RNAi increased the expression of HIF-1-responsive reporter. Transcription factor SKN-1 has been shown to have critical roles in enabling *C. elegans* to respond to oxidative stress [32–34, 39]. Our finding suggested a potential crosstalk between hypoxia response and oxidative stress response. To quantify the effect of *skn-1* RNAi on this HIF-1-responsive reporter, we examined *Pnhr-57:GFP* levels using protein blots. In the normal room air culture conditions, *Pnhr-57:GFP* was 40% higher in *skn-1* RNAi compared to control RNAi (Fig 2A) (*\*\*p* < 0.01, from three independent experiments). To gain insight to the effects of this interaction in hypoxic conditions, we moved the animals to 0.5% oxygen. After 4 hours of hypoxia treatment, *Pnhr-57:GFP* was 50% higher in *skn-1* RNAi compared to control RNAi (Fig 2A) (\*\**p* < 0.01, from three independent experiments). Thus, *skn-1* RNAi increased *Pnhr-57:GFP* levels under normoxic and hypoxic conditions.

**Fig 2.**
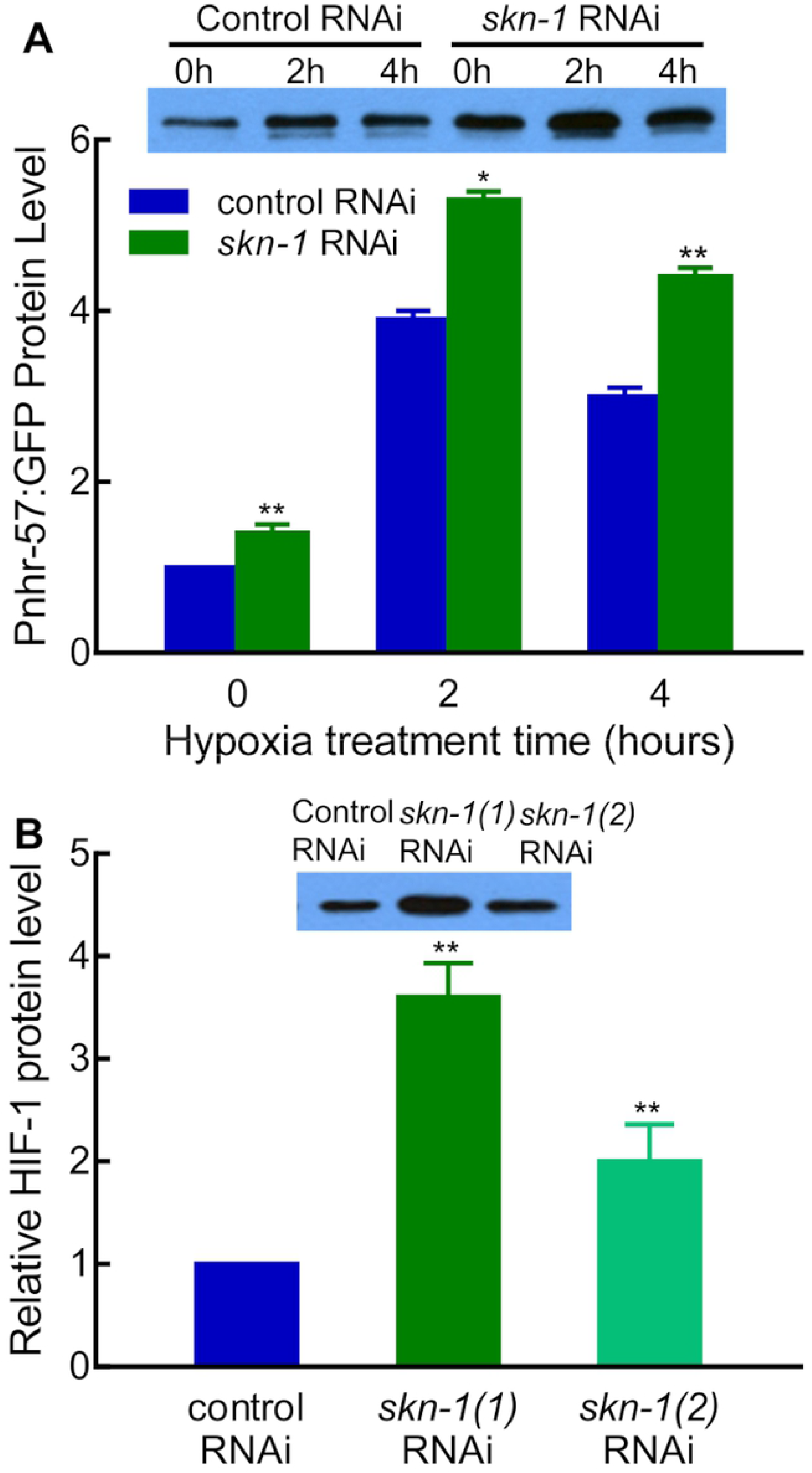
Identification of SKN-1 as a regulator of HIF-1. (A) *skn-1* RNAi increased expression of the *Pnhr-57:GFP* reporter. Reporter gene expression was quantitated in L4-stage animals in normal culture conditions or after 2 or 4 hours of hypoxia treatment (0.5% oxygen). The protein levels were calculated from three independent experiments and normalized to 0 hour hypoxia control RNAi. A representative western blot is shown. In each lane, lysates from 80 L4-satge worms were loaded. Asterisks indicate significant differences between control RNAi and *skn-1* RNAi at any given time point. *: *p* < 0.05; **:*p* < 0.01. (B) *skn-1* RNAi increased HIF-1 protein levels. These animals expressed an epitope-tagged HIF-1 protein [40]. The protein levels were calculated from three independent experiments and normalized to control RNAi. Two different RNAi clones were assayed, and they are designated as *skn-1(1)* and *skn-1(2)* here. A representative western blot is shown. In each lane, lysates from 100 L4-satge worms were loaded. Asterisks indicate significant differences between control RNAi and *skn-1* RNAi at any given time point. **: *p* < 0.01.

We next asked whether *skn-1* RNAi increased HIF-1 protein levels. We tested two *skn-1* RNAi constructs, and each resulted in an increase of HIF-1 protein levels by 2 to 3-fold as shown in Fig 2B (\*\**p* < 0.01, from three independent experiments). In sum, these results showed that *skn-1* RNAi increased HIF-1 protein level and HIF-1 reporter expression.

### Differential requirements for *skn-1* and *hif-1*

The finding that SKN-1 repressed HIF-1 protein levels suggested that there might be conditions in which it would be beneficial for the animal to express one of these two stress-responsive transcription factors, but not the other. To address this, we examined the relative requirements for *skn-1* and *hif-1* more closely.

In previous studies, we and others had shown that *hif-1* was required for survival in moderate hypoxia [25, 41]. As validated in the experiments described in Table 1, loss of *hif-1* impaired animal development and survival in 0.5% oxygen: after 24 hours of hypoxia treatment, only 75.8% of *hif-1*-deficient eggs hatched, and only 25.6% developed to adulthood within 72 hours. In contrast, *skn-1* RNAi had no effect on *C. elegans* development and survival in hypoxic conditions: after 24 hours of hypoxia treatment, 99.4% of *skn-1* RNAi treated eggs hatched and completed normal development to adulthood within 72 hours.

**Table 1.**
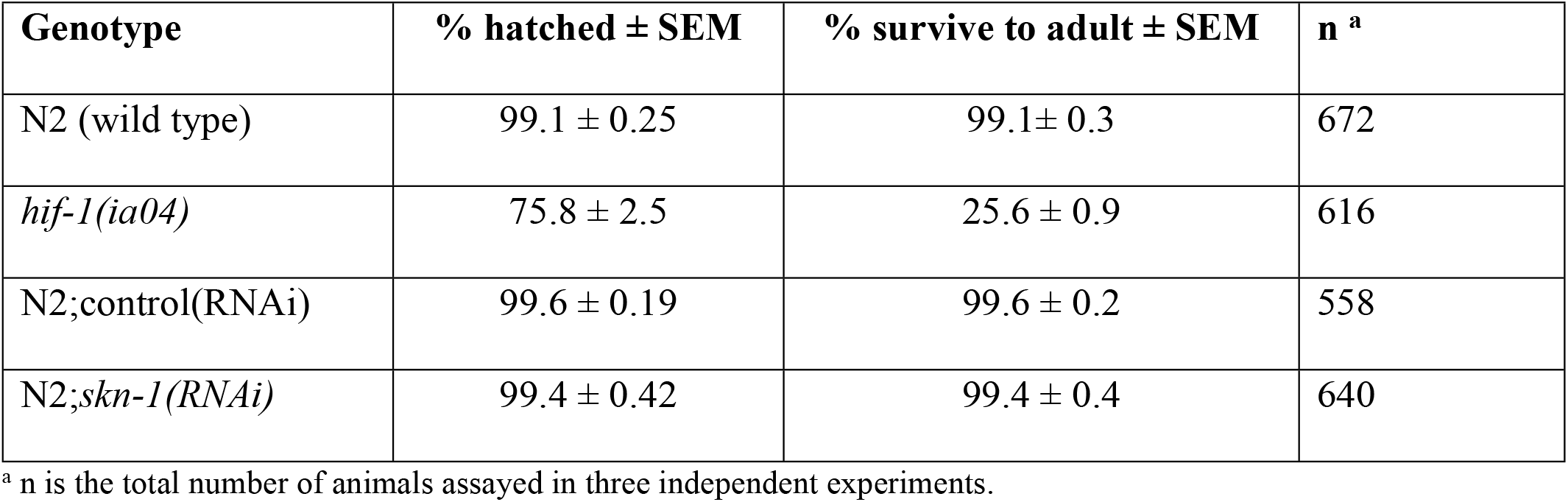
Relative requirements for *skn-1* and *hif-1:* survival in 0.5% oxygen.

| Genotype | % hatched ± SEM | % survive to adult ± SEM | n <sup>a</sup> |
| --- | --- | --- | --- |
| N2 (wild type) | 99.1 ± 0.25 | 99.1 ± 0.3 | 672 |
| *hif-1(ia04)* | 75.8 ± 2.5 | 25.6 ± 0.9 | 616 |
| N2;control(RNAi) | 99.6 ± 0.19 | 99.6 ± 0.2 | 558 |
| N2;*skn-1*(RNAi) | 99.4 ± 0.42 | 99.4 ± 0.4 | 640 |

<sup>a</sup> n is the total number of animals assayed in three independent experiments.

Prior studies also suggested that there were differential requirements for HIF-1 and SKN-1 in oxidative stress conditions. Mutants carrying loss-of-function mutations in *skn-1* have been shown to decrease the ability of *C. elegans* to survive exposure to agents that cause oxidative stress [34, 42–44]. In contrast, *C. elegans* carrying loss-of-function mutations in *hif-1* have been reported to be relatively resistant to peroxide [40]. We compared these phenotypes directly, and the data are provided in Table 2. These experiments confirmed that, while *skn-1*-deficient animals were sensitive to t-butyl peroxide, mutants lacking *hif-1* were remarkably resistant to this oxidative stress: while none of the *skn-1-*deficient mutants survived 6 hours of t-butyl peroxide treatment, 97.5% of *hif-1*-deficient mutants survived 10 hours of t-butyl peroxide treatment.

**Table 2.** Relative requirements for *skn-1* and *hif-1:* survival on t-butyl-peroxide.

| Exposure time | Genotype | Mean Survival ± SEM | N <sup>a</sup> |
| --- | --- | --- | --- |
| 6 hours | N2 (wild type) | 95.2 ± 2.6 | 106 |
| | *skn-1(zu67)* | 0.0 ± 0 | 117 |
| | *hif-1(ia04)* | 100.0 ± 0 | 120 |
| 8 hours | N2 (wild type) | 29.08 ± 7.7 | 106 |
| | *skn-1(zu67)* | 0.0 ± 0 | 117 |
| | *hif-1(ia04)* | 99.17 ± 0.8 | 120 |
| 10 hours | N2 (wild type) | 2.78 ± 2.8 | 106 |
| | *skn-1(zu67)* | 0.0 ± 0 | 117 |
| | *hif-1(ia04)* | 97.50 ± 2.5 | 120 |

<sup>a</sup> n is the total number of animals assayed in three independent experiments.

### SKN-1/NRF promotes *egl-9* expression

We next sought to discover the mechanism by which SKN-1 inhibits HIF-1 protein levels. In silico analyses identified a potential SKN-1 binding site in the *egl-9* promoter region (Fig 3A). EGL-9 is a central inhibitor of HIF-1 protein levels [26, 27] and of HIF-1 transcriptional activity [29, 30]. This suggested a model in which the SKN-1 DNA-binding complex bound directly to the *egl-9* regulatory sequences to promote *egl-9* expression, which, in turn, would ultimately decrease HIF-1 protein levels. To test this, we employed real-time quantitative RT-PCR to compare *egl-9* mRNA levels in worms fed with *skn-1* RNAi versus control RNAi. To produce reliable and reproducible results, *egl-9* mRNA levels were quantitated in three independent real-time quantitative RT-PCR experiments in L4-stage animals in room air or hypoxic conditions (0.5% oxygen). Each sample was performed with three technical replicates, and they produced similar C_t_ values. There are seven isoforms of *egl-9* mRNA transcripts (https://wormbase.org/species/c_elegans/gene/WBGene00001178#0-9f-10). The real-time quantitative PCR primer set used in this study can detect six *egl-9* mRNA isoforms. In room air, *skn-1* RNAi decreased *egl-9* mRNA levels by 30% compared to control RNAi (\*\**p* < 0.01, from three independent experiments) (Fig 3B). HIF-1 has been shown to activate *egl-9* mRNA expression under hypoxia, creating a negative feedback loop [27, 28]. In accordance with this, the inhibition effects of *skn-1* RNAi on *egl-9* mRNA levels were minimized by placing the animals in hypoxic conditions (Fig 3B).

**Fig 3.**
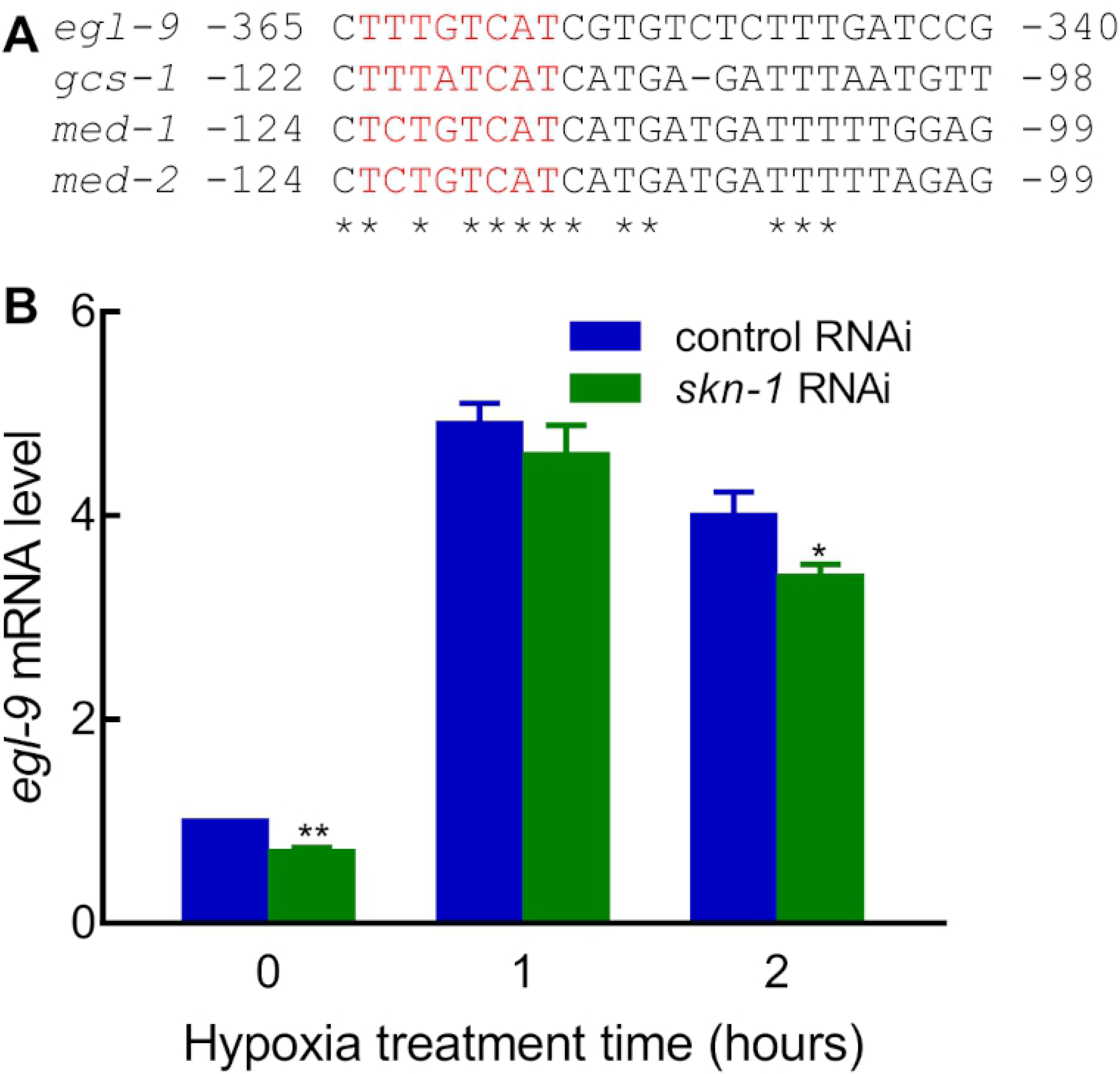
Identification of *egl-9* as a potential transcriptional target of SKN-1. (A) Sequence from the *egl-9* promoter was aligned with established SKN-1 binding sites in *gcs-1, med-1,* and *med-2.* Asterisks identify sequence identities shared by all four promoter regions in this interval, and predicted SKN-1 binding sites are in red. (B) *skn-1* RNAi decreased *egl-9* mRNA levels. *egl-9* mRNA levels were quantitated from three independent real-time quantitative RT-PCR experiments. The values at each time point were normalized to the 0 hour hypoxia control RNAi. Asterisks indicate significant differences between control RNAi and *skn-1* RNAi at any given time point. *:*p* < 0.05; **: *p* < 0.01.

To test the hypothesis that conditions that activate SKN-1 can promote *egl-9* promoter activity, we generated a reporter construct in which 1.6 kb of *egl-9* regulatory sequence directed the expression of GFP (Fig 4A). To distinguish the effects of SKN-1 on *egl-9* expression from those of HIF-1, we conducted these experiments in a *hif-1* mutant background. In agreement with prior studies [45], *Pegl-9:GFP* was visible in several tissues, including the body muscle, vulva, pharynx, anterior intestine, rectal cells and additional cells in the tail in standard culture conditions (20 °C) (Figs 4B and 4C). When the animals were treated with heat shock conditions that had been shown to activate SKN-1 (29 °C for 20 hours) [34], we observed dramatic induction of *Pegl-9:GFP* in the intestine. (Figs 4D and 4E).

**Fig 4.**
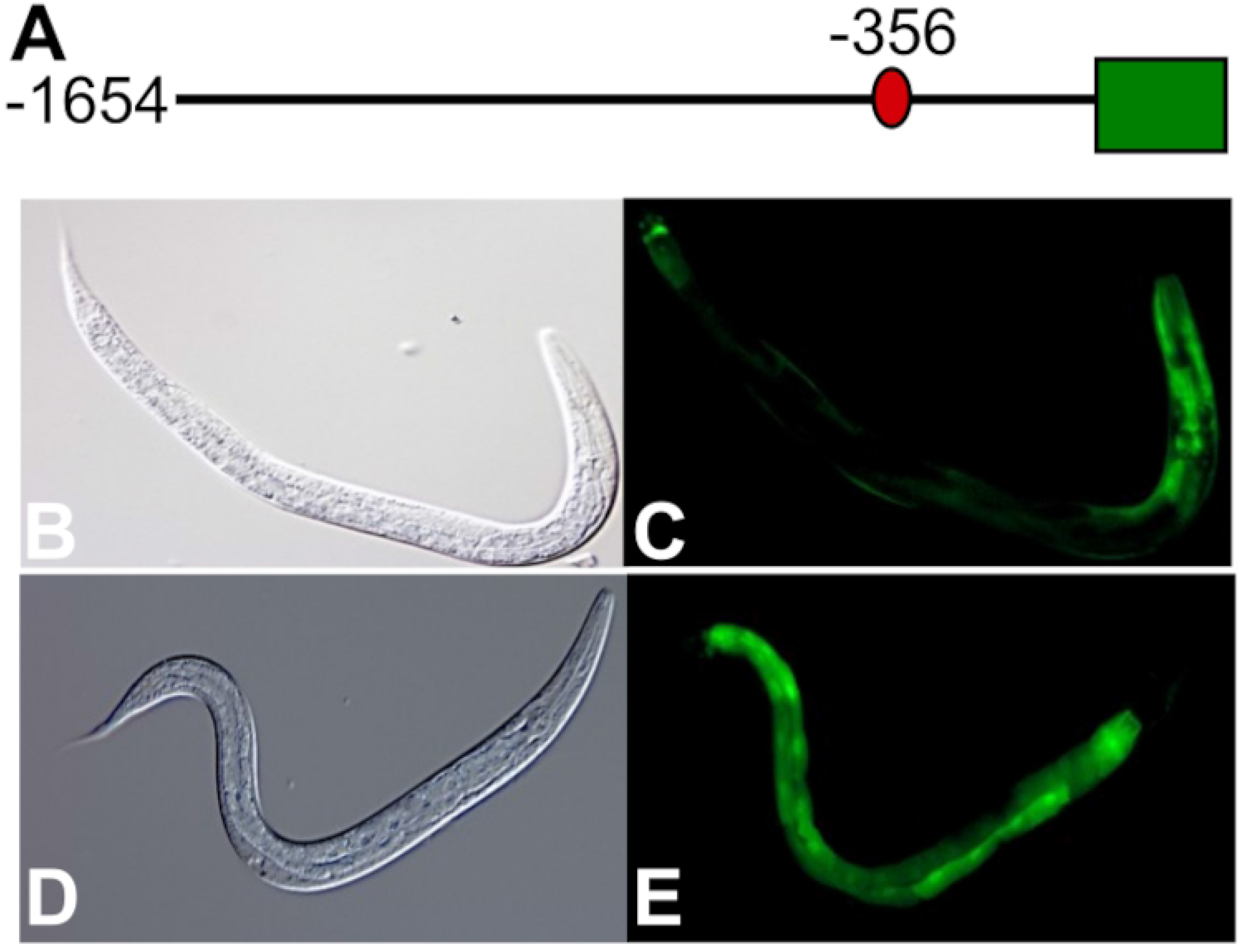
Heat shock alters *Pegl-9:GFP* expression. (A) The *Pegl-9:GFP* construct includes 1.6 kb of sequence 5’ to the *egl-9* translational start. GFP coding sequence is diagramed as a green box. The red oval indicates the position of the putative SKN-1 binding site. (B-E) *Pegl-9:GFP* expression in L4-stage animals under normal culture conditions and heat shock. Animals are shown as DIC images (B and D) and corresponding images of GFP fluorescence (C and E). In all images, the head is to the right. (B and C) Under normal conditions, *Pegl-9:GFP* was expressed in the body muscle, vulva, pharynx, anterior intestine, rectal cells and additional cells in the tail. (D and E) After heat shock treatment (29°C for 20 hours), *Pegl-9:GFP* was strongly induced in the intestine. *ThePegl-9:GFP* expression patterns in the L1, L2, L3 and adults were similar to that in the L4 worms, under both normal and heat shock conditions (data not shown).

We next asked whether heat shock induction of *Pegl-9:GFP* required *skn-1* function. We found that heat shock increased *Pegl-9:GFP* by 2.5-fold in animals carrying the wild-type *skn-1* allele. However, the heat shock induction of *Pegl-9:GFP* was abolished in *skn-1(zu67)* loss-of-function mutants (Fig 5A). Analyses of another independent *Pegl-9:GFP* transgenic line yielded similar results (data not shown).

To test the hypothesis that the putative SKN-1 binding site in the *egl-9* promoter was required for heat shock induction of *Pegl-9:GFP,* we generated the *P(m)egl-9:GFP* construct, which contained mutations in the putative SKN-1 binding site (in red type in Fig 3A). Heat shock increased *Pegl-9:GFP* by 2.1-fold. However, heat shock failed to induce the expression of *P(m)egl-9:GFP* (Fig 5B). Experiments with a second *P(m)egl-9:GFP* transgenic line gave similar results (data not shown). Collectively, these data demonstrated that heat shock induction of *Pegl-9:GFP* required *skn-1* function and the putative SKN-1 binding site in the *egl-9* promoter.

**Fig 5.**
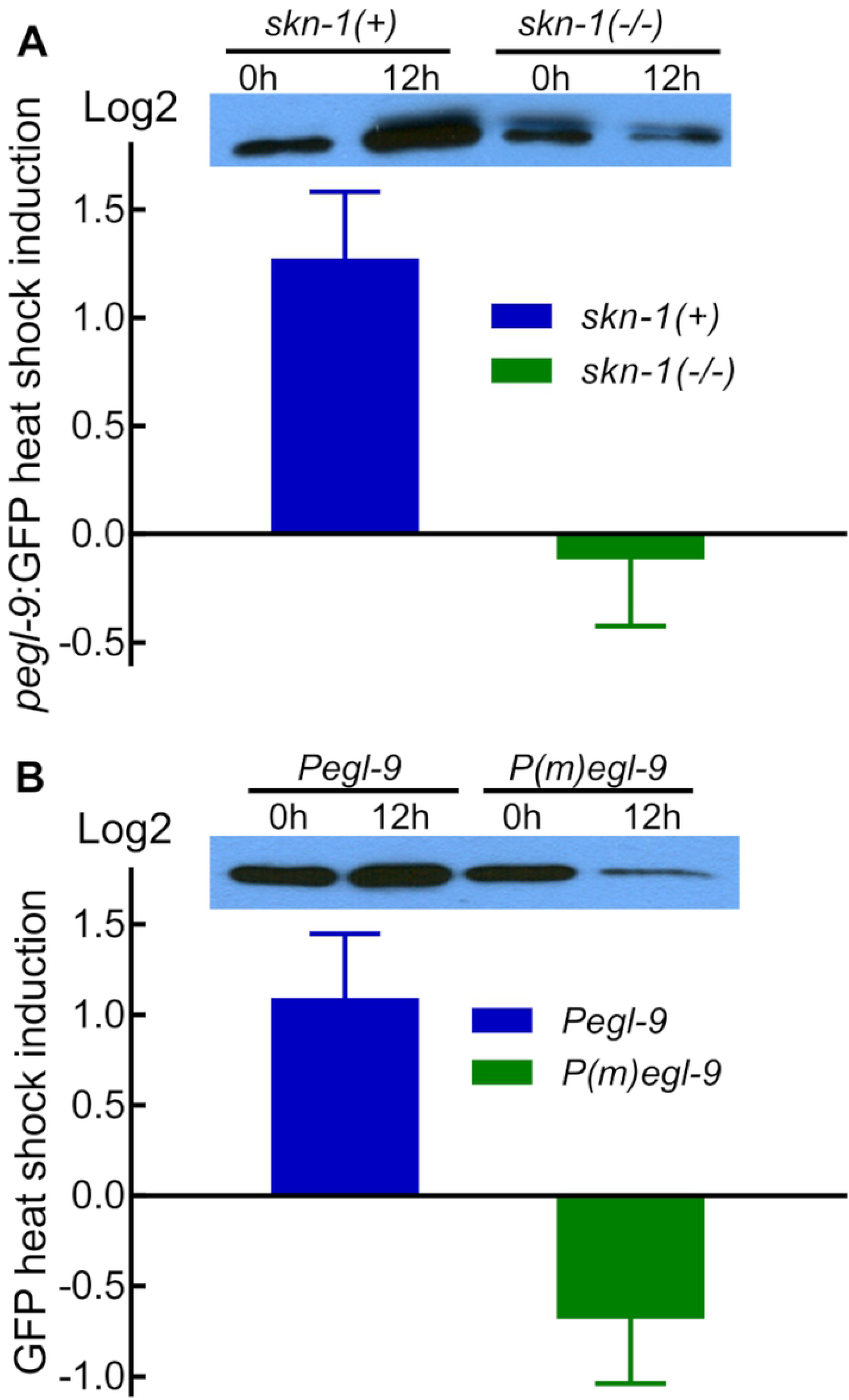
SKN-1 acts through the putative SKN-1 binding site in the *egl-9* promoter to activate *egl-9* expression. (A) Heat shock induced *Pegl-9:GFP* in animals carrying the wild-type *skn-1* allele, but did not induce the reporter in animals carrying the *skn-1(zu67)* loss-of-function mutation. The vertical axis shows the log2 fold changes of GFP caused by heat shock for each strain with standard errors, as determined by four biological replicates. A representative western blot is shown. For each sample, 20 L4-satge worms were boiled and lysates corresponding to 10 worms were loaded to each lane. (B) Heat shock increased the expression of *Pegl-9:GFP,* but did not increase the expression of the reporter in which the putative SKN-1 binding site was mutated (*P(m)egl-9:GFP)*. The vertical axis shows the log2 fold changes of GFP caused by heat shock for each strain with standard errors, as determined by five biological replicates. A representative western blot is shown. For each sample, 20 L4-satge worms were boiled and lysates corresponding to 10 worms were loaded to each lane.

## Discussion

This study provides new insights to the mechanisms that allow animals to respond appropriately to diverse stresses. While HIF-1 and SKN-1 are both stress responsive transcription factors, they have distinct functions. For example, *skn-1*-deficient animals are less able to survive exposure to peroxide, while *hif-1-deficient* mutants are relatively resistant to this oxidizing agent [40]. Conversely, while a deletion mutation in *hif-1* dramatically impairs survival in 0.5% oxygen [25], *skn-1* RNAi has little effect (Table 1). Thus, animals may benefit from cross-talk between these two transcription factors as they are challenged by oxygen deprivation and oxidative stress. Here, we provide evidence that SKN-1 promotes *egl-9* expression, thereby attenuating HIF-1 function.

### SKN-1 promotes *egl-9* expression, thereby inhibiting HIF-1

Our data support a model in which SKN-1 binds directly to the *egl-9* promoter to increase *egl-9* expression. This conclusion is further substantiated by the genome wide chromatin immunoprecipitation experiments that determined that SKN-1:GFP associated with DNA sequences in the *egl-9* 5’ regulatory region [46]. EGL-9 functions as a cellular oxygen sensor, and it mediates oxygen-dependent degradation of HIF-1 [26]. Hence, *skn-1* RNAi results in increased HIF-1 protein levels (Fig 2B). We expect that this regulatory interaction, in which SKN-1 can quickly down-regulate HIF-1, would allow animals to adapt quickly to changes in cellular conditions.

Prior studies have identified genes that are positively regulated by either SKN-1 or HIF-1, and these gene lists are largely non-overlapping [28, 32, 47–49]. Genes that are commonly regulated by both SKN-1 and HIF-1 include K10H10.2/*cysl-2,* F57B9.1, M05D6.5, and *rhy-1.* Two lines of evidence suggest that *rhy-1* may be a direct target of SKN-1. First, Oliveira et al. (2009) identified a potential SKN-1 binding site in the *rhy-1* promoter region. Second, the modENCODE project found that *rhy-1* 5’ regulatory sequences associated with SKN-1 *in vivo* [46]. Like *egl-9, rhy-1* is also a negative regulator of HIF-1 [28, 29]. Collectively, these data suggest that SKN-1 may act through both *egl-9* and *rhy-1* to reduce HIF-1 function. Interactions between HIF-1 and SKN-1 are likely to be different in select cells, developmental stages, or environmental contexts. HIF-1, EGL-9 and SKN-1 each have other developmental functions, and some of these are specific to certain cell types [34, 50–54]. The downstream effects of SKN-1 or HIF-1 activation are further influenced by cellular or environmental contexts. For example, *C. elegans* SKN-1 can be activated by arsenite or by t-butyl hydroperoxide, but only a subset of SKN-1 targets are activated by either toxicant [32]. Also, while *C. elegans* SKN-1 and HIF-1 have distinct roles in peroxide and hypoxia stress responses, their functions overlap in selenium and hydrogen sulfide stress responses [13, 15, 35–37]. Similarly, the mammalian NRF and HIF transcription factors are very sensitive to environmental and physiological cues or stresses, and their regulatory relationships are context specific. Ischemia causes tissue hypoxia, which stabilizes HIF transcription factors. Once the ischemic tissue is reperfused, HIF transcription factors are degraded quickly and NRF2 is up-regulated, presumably to limit the oxidative damage [4, 5]. However, intermittent hypoxia has been shown to induce both HIF-1α and NRF2 [6, 7]. While NRF2 signaling activates HIF-1 in several cancer types [12], studies of the anti-inflammatory drug andrographolide in endothelial cells revealed interactions between NRF2 and the PHDs that modulate HIF-1 [55]. NRF2 and the HIF transcription factors have key roles in angiogenesis and iron regulation, and their functions can converge on developmental processes or feedback loops that modulate their activities [12, 56–58].

### *Pnhr-57:GFP* as a marker for hypoxia-induced gene expression: discoveries and insights from a genome-wide screen

Prior studies have demonstrated that *nhr-57* was induced by hypoxia in a *hif-1*-dependent manner and that over-expression of *Pnhr-57:GFP* in *egl-9* mutants required *hif-1* [13, 27–30]. Moreover, Bellier *et al.* (2009) found that HIF-1-mediated induction of *nhr-57* helped to protect *C. elegans* from the lethal effects of pore-forming toxins [59]. While these studies showed that *nhr-57* is a direct target of HIF-1, other transcription factors must also contribute to its expression. The studies presented here show that, while genes such as *egl-9* or *rhy-1* clearly regulate HIF-1 to control *Pnhr-57:GFP* expression, many other RNAi treatments can activate *Pnhr-57:GFP* through *hif-l*-independent pathways. The *Pnhr-57:GFP* reporter will continue to be a valuable marker, but these data inform our interpretations of studies that employ this reporter. While *Pnhr-57:GFP* is clearly regulated by HIF-1, it is important to compare expression of the reporter in wild-type and *hif-1*-deficient animals before drawing conclusions about HIF-1 activity.

We found that diverse RNAi treatments that compromise metabolic function or protein homeostasis increased *Pnhr-57:GFP* expression, and this effect did not require a functional *hif-1* gene. Interestingly, many of the genes integral to these processes have been shown to have roles in stress response and aging [60–65]. RNAi-mediated depletion of proteasomal components has also been shown to impact resistance to polyglutamine toxicity and to induce expression of *Pgpdh-1:GFP,* a marker for osmotic stress and glycerol production [66, 67].

Further characterization of the 179 RNAi treatments that increased *Pnhr-57*:GFP identified 13 genes that had much stronger effects in animals carrying a wild-type *hif-1* gene. These genes included *vhl-1, egl-9,* and *rhy-1.* These three genes had all been identified in prior studies as negative regulators of HIF-1 [26, 29]. The succinate dehydrogenase subunit *sdhb-1* was also found to have *hif-1*-dependent effects. This is especially interesting, since studies in cancer cell lines have shown that succinate can inhibit the enzymatic activities of HIF prolyl hydroxylases [68, 69]. *sams-1* and *sbp-1* encode the *C. elegans* S-adenosyl methionine synthetase and the SREBP homologs, respectively. The RNAi treatments of sams-1 and sbp-1 have a lesser impact on *Pnhr-57:GFP* levels in *hif-1* mutants, suggesting that the effects of *sams-1* and *sbp-1* RNAi on the reporter are mediated by HIF-1 (S2 Fig). Both of these genes have key roles in methionine metabolism and fatty acid biosynthesis [70], and it will be interesting to investigate the ways in which these important processes intersect with hypoxia response.

### Materials and methods

#### Strains

The following strains were used in this study: wild-type N2 Bristol; ZG430: *Pnhr-57:GFP(iaIs07)TV; egl-9(sa307)V; hif-1(ia04)V; Phif-1:hif-1a:Myc:HA (iaIs28);* ZG120: *Pnhr-57:GFP(iaIs07)IV;* ZG509: *rrf-3(pk1426)II; Pnhr-57:GFP(iaIs07)IV;* ZG508: *rrf-3(pk1426)II; Pnhr-57:GFP(iaIs07)IV; hif-1(ia04)V;* ZG429: *hif-1(ia04)V; Phif-1:hif-1a:Myc:HA(ials28);* ZG472: *hif-1(ia04)V; Pegl-9:GFP(iaEx84);* ZG487: *hif-1(ia04)V; P(m)egl-9:GFP(iaEx96);* ZG488: *skn-1(zu67)IV; hif-1(ia04)V; Pegl-9:GFP(iaEx84).* The transgenes expressing epitope-tagged HIF-1 protein were described and characterized previously [40].

#### RNAi experiments

The RNAi screen was conducted as previously described [71], with few modifications. Each bacterial clone (expressing double-stranded RNA for one gene) was cultured in L-broth with 50 ug/mL ampicillin and 12.5 ug/mL tetracycline overnight at 37°C. The following morning, the bacteria were inoculated into new L-broth with 100 ug/mL ampicillin for 6 hours at 37°C before seeding on 24-well NGM agar plates with 25 ug/mL carbenicillin and 2 mM IPTG. Each RNAi clone was plated in duplicate. The following day, 15-25 L1-stage worms were added to each well. The plates were incubated at 15°C for 5-6 days, and then the worms were screened for positive *Pnhr-57:GFP* green fluorescence by stereomicroscopy. Bacterial RNAi clones that increased the reporter were rescreened in two independent replicates, and the plasmid inserts were validated by sequencing.

*skn-1* RNAi causes maternal-effect lethality, in this study we examined the effects of first generation *skn-1* RNAi. Dead egg percentages given by the first generation *skn-1* RNAi adults were measured to check the *skn-1* RNAi efficiency. We routinely achieved as high as 90% dead egg percentages from the first generation *skn-1* RNAi adults, indicating high *skn-1* RNAi efficiency.

#### Hypoxia stress and oxidative stress assays

To assess the relative effects of t-butyl-peroxide exposure, animals in the first day of adulthood were placed on NGM plates containing 7.5 mM t-butyl-peroxide, in the presence of bacterial food. The survival was scored after treating the animals for 6, 8 or 10 hours.

For hypoxia experiments, adults were allowed to lay eggs on standard NGM plates with OP50 bacterial food for 2 hours. The adults were then removed, and the plates with embryos were placed in a hypoxia chamber (0.5% oxygen; 21°C) for 24 hours. After 24 hours, the plates were removed from the hypoxia chamber, and the un-hatched eggs were counted immediately. The plates were then maintained in room air (21°C). The adult worms were counted at 72 hour since the eggs had been laid. Wild-type control animals hatched within 24 hours and reached adulthood within 72 hours.

#### *Pegl-9:GFP* expression constructs

To generate the *Pegl-9:GFP* construct, a fragment that contained 1.6 kb of sequence upstream of the initiation ATG of *egl-9* gene was amplified by PCR using the forward primer 5’-CGCGCATGCGTGTATGTGTGTGAAAGAG-3’ and the reverse primer 5’-GCGGTCGACGCAACTTTTTTCTGTCACATTCAG-3’. The PCR product was cloned into the green fluorescence protein (GFP) vector pPD95.75 (gift from Andrew Fire). To create the *P(m)egl-9:GFP* point mutation construct, the predicted SKN-1 binding site TTTGTCAT [34, 72]was altered to CGACGGGC. Transgenic animals were generated by injection of DNA into the gonadal syncitium, using standard methods with *rol-6* (pRF4) as the co-injection marker [73]. For each construct, two independent transgenic lines were generated and assayed.

#### Protein Blots

To assay the expression of GFP or HIF-1 proteins, 20-100 L4-stage worms were collected and boiled for 5 min in 1X SDS sample buffer, and the lysates were size fractionated on polyacrylamide gels and analyzed by Western blots. The GFP-specific mouse monoclonal antibody (from Roche) was used at 1:500. The HA-specific mouse monoclonal antibody (from Cell Signaling) was used at 1:250. The secondary antibody (goat anti-mouse IgG+IgM from Biorad) was used at 1:2000 dilutions. The western blot images were analyzed by the Image J software. For each assay, three to five independent biological replicates were included.

#### RNA extraction and real-time quantitative RT-PCR

Total RNA was isolated from synchronized L4-stage animals using Trizol (Invitrogen) and RNeasy Mini Kit (Qiagen). After being treated by RNase free DNase (Promega), total RNA was reverse transcribed to complementary DNA using Oligo dT_18_ primer and AffinityScript reverse transcriptase (Stratagene). Real-time quantitative PCR was performed using the iQ SYBR GREEN supermix (Bio-Rad) and Stratagene Mx4000 multiplex PCR system. In the assay, three biological replicates were included. And for each sample, three technical replicates were performed and they gave similar C_t_ values. And cDNA from 62.5 ng of total RNA was added to each PCR reaction. Relative mRNA quantification was performed using the efficiency-corrected comparative quantification method [74]. *inf-1*, a gene not regulated by hypoxia, was used as the reference gene [28]. The primer sequences for *inf-1* real-time quantitative PCR are included in S1 Fig. The primers for *egl-9* real-time quantitative PCR are the forward primer 5’-GCCGACTTTCAATCCACTTC-3’ and reverse primer 5’-AATGATCGGAGATCGACTGG-3’. There are seven isoforms of *egl-9* mRNA transcripts (https://wormbase.org/species/c_elegans/gene/WBGene00001178#0-9f-10). This primer set can detect six out of seven *egl-9* mRNA isoforms. The isoform d.1 will not be detected by this primer set, because the forward primer is located within the eighth intron of the unspliced isoform d.1.

## Supporting information

**S1 Fig. *nhr-57* promoter HIF-1 chromatin immunoprecipitation experiments.** (A) *nhr-57* promoter sequence and positions of primers used for real-time quantitative PCR assays in HIF-1 chromatin immunoprecipitation experiments. (B) Primer sequences for real-time quantitative PCR assays in *nhr-57* promoter HIF-1 chromatin immunoprecipitation experiments. (C) Chromatin co-immunoprecipitation data. In these experiments the endogenous *hif-1* locus was disrupted by the *ia04* large deletion and HIF-1 function was restored by the *Phif-1:hif-1a:Myc:HA* transgene [40]. The relative amounts of *nhr-57* promoter regions that co-immunoprecipitated with HIF-1:Myc:HA was determined by real-time quantitative PCR. The bars show the average enriched fold from at least three independent replicates. *inf-1*, a gene not regulated by HIF-1, was used as the reference gene.

**S2 Fig. RNAi inactivation of *sams-1* or *sbp-1* increased *Pnhr-57:GFP* expression.** (A) RNAi for *sams-1* (S-adenosyl methionine synthetase) increased *Pnhr-57:GFP* expression more than 7-fold in animals carrying the wild-type *hif-1* allele relative to control RNAi, and increased the reporter 3-fold in animals carrying the *hif-1(ia04)* deletion. The difference in RNAi effect between *hif-1(+)* and *hif-1(ia04)* strains is statistically significant (*\*p* < 0.05, from six independent experiments, by student *t*-test). (B) RNAi for the SREBP homolog *sbp-1* increased expression of the reporter more than 3-fold in animals carrying the wild-type *hif-1* allele, but had no effect on *Pnhr-57:GFP* expression in *hif-1(ia04)* mutants. The difference in RNAi effect between *hif-1(+)* and *hif-1(ia04)* strains is statistically significant (\**p* < 0.05, from five independent experiments, by student *t*-test). GFP levels were determined by protein blots, and the control animals were fed on bacteria carrying the empty RNAi vector (L4440). The experiments were conducted in RNAi-sensitive strains (*rrf-3(pk1426))*.

**S1 Table. Genes increased *Pnhr-57:GFP* expression when knocked-down by RNAi.**

**S2 Table. Genes for which RNAi caused *hif-1*-dependent increase of *Pnhr-57:GFP* expression.**

