## Supplement Figure 1 for "Crosstalk in oxygen homeostasis networks: SKN-1/NRF inhibits the HIF-1 hypoxia-inducible factor in *Caenorhabditis elegans*"

**S1 Fig. *nhr-57* promoter HIF-1 chromatin immunoprecipitation experiments.**

(A) *nhr-57* promoter sequence and positions of primers used for real-time quantitative PCR assays in HIF-1 chromatin immunoprecipitation experiments.

1F 🡪

ttgaataatttgattagctgattggcttatgacatcagctgcgggaagtcattatgcaaacgtgaataatttggtagtacgtgtgttttattgttttctattttttgaattcaataacttgaataagaaaaatataattgaaaactaacaatgatgcaaacaaacaaaata

2F 🡪	🡨1R

aacttacatcatatctttcgcttcttcaactcaaagagtgtttattgtactttacttttctctctctccctctcactctccatacctttttcatttcccaagtgtctatcaacaaaacatacacaaaaagaggcgtaactctcactcaaaagatcagctggagaactatgcgtacgtgattagtggaacgtctaacctcccgcgtctccacattcaatcgatcactat

3F🡪	🡨2R

ccgaacaaattgtcatcgttttcggaaaacccagacctcgctgtaattttttgcatgatttccatctttgagacaccattttggtttgcatttctcacgtagacttcctggaggtcaccggtcttttatcttatcaatcgtaattgttagactatataaaacgtgcggtgt

🡨3R

tcaactacatgcttttgtatttttaaagagtgaatcaagag**atg**ttggtggctcgcg

(B) Primer sequences for real-time quantitative PCR assays in *nhr-57* promoter HIF-1 chromatin immunoprecipitation experiments.

*nhr-57*_1F 5’-attggcttatgacatcagctgc-3’

*nhr-57*_1R 5’-cactctttgagttgaagaagcg-3’

*nhr-57*_2F 5’-cgcttcttcaactcaaagagtg-3’

*nhr-57*_2R 5’-ttacagcgaggtctgggttttc-3’

*nhr-57*_3F 5’-gaaaacccagacctcgctgtaa-3’

*nhr-57*_3R 5’-ccaacatctcttgattcactct-3’

*inf-1* F 5’-ttccgttcaggatcttcccgtgtt-3’

*inf-1* R 5’-tctcggtgacgaagttgatggcaa-3’

(C) Chromatin co-immunoprecipitation data.

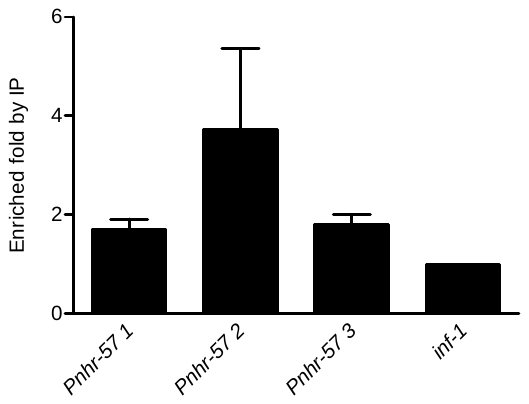

In these experiments the endogenous *hif-1* locus was disrupted by the *ia04* large deletion and HIF-1 function was restored by the *Phif-1:hif-1a:Myc:HA* transgene [1]. The relative amounts of *nhr-57* promoter regions that co-immunoprecipitated with HIF-1:Myc:HA was determined by real-time quantitative PCR. The bars show the average enriched fold from at least three independent replicates. *inf-1*, a gene not regulated by HIF-1, was used as the reference gene.

1.	Zhang Y, Shao Z, Zhai Z, Shen C, Powell-Coffman JA. The HIF-1 hypoxia-inducible factor modulates lifespan in C. elegans. PLoS One. 2009;4(7):e6348. Epub 2009/07/28. doi: 10.1371/journal.pone.0006348. PubMed PMID: 19633713.
