## Supplement Table 1 for "Crosstalk in oxygen homeostasis networks: SKN-1/NRF inhibits the HIF-1 hypoxia-inducible factor in *Caenorhabditis elegans*"

**S1 Table. Genes increased *Pnhr-57:GFP* expression when knocked-down by RNAi.**

| GenePair | Gene name | Predicted function | Functional category | [Identified Hansen M *et al.,* 2005](http://wormbase.org/db/misc/person?name=WBPerson2242;class=Person) | Identified in Hamilton B *et al.,* 2005 | Identified in Curran and Ruvkun 2007 | Identified in Nollen EAA *et al.,* 2004 | Identified in Kim Y and Sun H 2007 |
| --- | --- | --- | --- | --- | --- | --- | --- | --- |
| ZK809.3 |  | NADH:ubiquinone oxidoreductase | Metabolism: ETC |  |  |  |  | paraquat resistance; |
| F53F4.10 |  | NADH:ubiquinone oxidoreductase, NDUFV2 | Metabolism: ETC |  |  |  |  |  |
| K04G7.4 | nuo-4 | NADH:ubiquinone oxidoreductase NDUFA10 | Metabolism: ETC | [lifespan extended](http://wormbase.org/db/misc/person?name=WBPerson2242;class=Person) | lifespan extended |  |  | paraquat resistance; life span extended |
| Y63D3A.7 |  | NADH-ubiquinone oxidoreductase NDUFA2/B8 | Metabolism: ETC |  |  |  |  |  |
| C34B2.8 |  | NADH:ubiquinone oxidoreductase, B16.6 subunit | Metabolism: ETC |  |  |  |  |  |
| F59C6.5 |  | NADH-ubiquinone oxidoreductase, NDUFB10 | Metabolism: ETC |  |  | fat content; sod-3:GFP; lifespan extended |  |  |
| C33A12.1 |  | NADH:ubiquinone oxidoreductase, NDUFA5/B13 | Metabolism: ETC |  |  |  |  | paraquat resistance; |
| C16A3.5 |  | NADH:ubiquinone oxidoreductase, NDUFB9/B22 | Metabolism: ETC |  |  |  |  |  |
| F37C12.3 |  | NADH-ubiquinone oxidoreductase NDUFAB1/SDAP | Metabolism: ETC |  |  |  |  |  |
| T20H4.5 |  | NADH:ubiquinone oxidoreductase, NDUFS8 | Metabolism: ETC |  | lifespan extended |  |  | paraquat resistance; life span extended |
| C18E9.4 |  | NADH:ubiquinone oxidoreductase, NDUFB3/B12 | Metabolism: ETC |  | lifespan extended |  |  |  |
| C09H10.3 | nuo-1 | NADH:ubiquinone oxidoreductase, NDUFV1 | Metabolism: ETC |  |  | fat content; sod-3:GFP; lifespan extended |  |  |
| [ZK973.10](http://www.wormbase.org/db/seq/sequence?name=ZK973.10;class=Gene_name) | lpd-5 | NADH:ubiquinone oxidoreductase, NDUFS4 | Metabolism: ETC |  |  |  |  |  |
| T10E9.7 | nuo-2 | NADH-ubiquinone oxidoreductase, NDUFS3 | Metabolism: ETC | [lifespan extended](http://wormbase.org/db/misc/person?name=WBPerson2242;class=Person) |  |  |  |  |
| W01A8.4 |  | NADH-ubiquinone oxidoreductase NDUFB4/B15 | Metabolism: ETC |  |  |  |  |  |
| F22D6.4 | nduf-6 | NADH:ubiquinone oxidoreductase, NDUFS6 | Metabolism: ETC |  |  |  |  |  |
| D2030.4 |  | NADH:ubiquinone oxidoreductase, NDUFB7/B18 | Metabolism: ETC |  | lifespan extended |  |  |  |
| Y57G11C.12 | nuo-3 | NADH:ubiquinone oxidoreductase, NDUFA6/B14 & BRICK1 | Metabolism: ETC | [lifespan extended](http://wormbase.org/db/misc/person?name=WBPerson2242;class=Person) |  |  |  | paraquat resistance; |
| T26A5.3 | nduf-2.2 | NADH:ubiquinone oxidoreductase, NDUFS2 | Metabolism: ETC |  |  |  |  | paraquat resistance; life span extended |
| W02F12.5 |  | oxoglutarate dehydrogenase complex / E2 subunit | Metabolism: Mitochondrial |  |  |  |  |  |
| T07C4.7 | mev-1 | succinate dehydrogenase complex / cytochrome b | Metabolism: ETC |  |  |  |  |  |
| F42A8.2 | sdhb-1 | succinate dehydrogenase complex, subunit B | Metabolism: ETC |  |  |  |  |  |
| F33A8.5 | sdhd-1 | succinate dehydrogenase complex, subunit D | Metabolism: ETC |  |  |  |  |  |
| F56D2.1 | ucr-1 | ubiquinol-cytochrome c oxidoreductase complex | Metabolism: ETC |  |  |  |  | paraquat resistance; life span extended |
| Y37D8A.14 | cco-2 | Cytochrome c oxidase, subunit Va/COX6 | Metabolism: ETC | [lifespan extended](http://wormbase.org/db/misc/person?name=WBPerson2242;class=Person) |  |  |  | paraquat resistance; life span extended |
| F45H10.2 |  | Ubiquinol cytochrome c reductase, subunit QCR8 | Metabolism: ETC |  |  |  |  |  |
| T02H6.11 | phi-44 | Ubiquinol cytochrome c reductase, subunit QCR7 | Metabolism: ETC |  |  |  | unc-54:POLYQ(Q35)YFP aggregation |  |
| C01F1.2 | sco-1 | putative cytochrome c oxidase assembly protein | Metabolism: ETC |  |  |  |  |  |
| C54G4.8 | cyc-1 | cytochrome c reductase component | Metabolism: ETC | [lifespan extended](http://wormbase.org/db/misc/person?name=WBPerson2242;class=Person) |  |  |  |  |
| F26E4.6 |  | Cytochrome c oxidase, subunit VIIc/COX8 | Metabolism: ETC |  | lifespan extended | fat content; sod-3:GFP; lifespan extended |  |  |
| F26E4.9 | cco-1 | Cytochrome c oxidase, subunit Vb/COX4 | Metabolism: ETC | [lifespan extended](http://wormbase.org/db/misc/person?name=WBPerson2242;class=Person) | lifespan extended |  |  |  |
| W09C5.8 |  | Cytochrome c oxidase, subunit IV/COX5b | Metabolism: ETC |  | lifespan extended |  |  |  |
| R04F11.2 |  | Mitochondrial F1F0-ATP synthase, subunit e | Metabolism: ETC |  |  |  |  |  |
| T05H4.12 | atp-4 | Mitochondrial F1F0-ATP synthase, subunit Cf6 | Metabolism: ETC | [lifespan extended](http://wormbase.org/db/misc/person?name=WBPerson2242;class=Person) |  |  |  |  |
| F02E8.1 | asb-2 | Mitochondrial F1F0-ATP synthase, subunit b/ATP4 | Metabolism: ETC | [lifespan extended](http://wormbase.org/db/misc/person?name=WBPerson2242;class=Person) |  |  | unc-54:POLYQ(Q35)YFP aggregation |  |
| C53B7.4 | asg-2 | Mitochondrial F1F0-ATP synthase, subunit g / ATP20 | Metabolism: ETC |  | lifespan extended |  |  |  |
| C34E10.6 | atp-2 | F0F1-type ATP synthase, beta subunit | Metabolism: ETC |  |  | fat content; sod-3:GFP; lifespan extended; polyQ agg. | unc-54:POLYQ(Q35)YFP aggregation |  |
| H28O16.1 | phi-37 | F0F1-type ATP synthase, alpha subunit | Metabolism: ETC |  |  | sod-3:GFP; lifespan extended; polyQ agg. |  |  |
| Y82E9BR.3 |  | Mitochondrial F1F0-ATP synthase, subunit c/ATP9 | Metabolism: ETC |  |  |  |  |  |
| R53.4 |  | Mitochondrial F1F0-ATP synthase, subunit f | Metabolism: ETC |  |  |  |  |  |
| F27C1.7 | atp-3 | Mitochondrial F1F0-ATP synthase, subunit OSCP/ATP5 | Metabolism: ETC | [lifespan extended](http://wormbase.org/db/misc/person?name=WBPerson2242;class=Person) |  | fat content; sod-3:GFP; lifespan extended; polyQ agg.; DAF-16:GFP loc. |  |  |
| K07A12.3 | asg-1 | Mitochondrial F1F0-ATP synthase, subunit g/ATP20 | Metabolism: ETC |  |  |  |  |  |
| F58F12.1 | phi-38 | Mitochondrial F1F0-ATP synthase, subunit delta/ATP16 | Metabolism: ETC |  |  |  | unc-54:POLYQ(Q35)YFP aggregation |  |
| B0261.4 |  | mitochondrial ribosomal protein L47 | Metabolism: Mito translation |  | lifespan extended |  |  |  |
| B0511.8 | tag-264 | Mitochondrial 28S ribosomal protein S30 | Metabolism: Mito translation |  |  |  |  |  |
| B0432.3 | mrpl-41 | Mitochondrial ribosomal protein L27 | Metabolism: Mito translation |  |  |  |  |  |
| T21B10.1 |  | uncharacterized protein, similarity to mitochondrial ribosomal subunit | Metabolism: Mito translation |  |  | lifespan extended |  |  |
| Y54E10A.7 |  | similar to mitochondrial 39S ribosomal protein L17 | Metabolism: Mito translation |  |  |  |  |  |
| F59A3.3 |  | similar to mitochondrial ribosomal protein L24 | Metabolism: Mito translation |  |  |  |  |  |
| Y119D3B.16 |  | similar to mitochondrial 39S protein L45 | Metabolism: Mito translation |  |  |  |  |  |
| W04D2.5 |  | Mitochondrial ribosomal protein S11 | Metabolism: Mito translation |  |  |  |  |  |
| E02A10.1 | mrps-5 | mitochondrial ribosomal protein S5 | Metabolism: Mito translation |  |  |  |  |  |
| T23B12.3 |  | Mitochondrial ribosomal protein S2 | Metabolism: Mito translation |  |  |  |  |  |
| T23B12.2 |  | Mitochondrial ribosomal protein L4 | Metabolism: Mito translation |  |  |  |  |  |
| F33D4.5 |  | mitochondrial 50S ribosomal protein L1 | Metabolism: Mito translation |  |  |  |  | paraquat resistance; |
| F56B3.8 |  | Mitochondrial ribosomal protein L2 | Metabolism: Mito translation |  |  |  |  |  |
| Y55F3BL.1 |  | Similar to mitochondrial ribosomal protein L46 | Metabolism: Mito translation |  |  |  |  | paraquat resistance; |
| W04B5.4 |  | Mitochondrial ribosomal protein L30 | Metabolism: Mito translation |  |  |  |  |  |
| T04A8.11 |  | Mitochondrial ribosomal protein L16 | Metabolism: Mito translation |  |  |  |  |  |
| C05D11.10 |  | Mitochondrial ribosomal S17-like protein | Metabolism: Mito translation |  |  |  |  |  |
| C26E6.6 |  | similar to mitochondrial 39S ribosomal protein L3 | Metabolism: Mito translation |  |  |  |  |  |
| [CD4.3](http://www.wormbase.org/db/seq/sequence?name=CD4.3;class=Gene_name) |  | similar to 39S ribosomal protein L48 | Metabolism: Mito translation |  |  |  |  |  |
| F21D5.8 |  | putative mitochondrial 28S ribosomal protein | Metabolism: Mito translation |  |  |  |  | paraquat resistance; |
| F09G8.3 |  | Mitochondrial ribosomal protein S9 | Metabolism: Mito translation |  |  |  |  | paraquat resistance; life span extended |
| F29B9.10 |  | mitochondrial 28S ribosomal protein S21 | Metabolism: Mito translation |  |  |  |  |  |
| F46B6.6 |  | Mitochondrial translation initiation factor 2 | Metabolism: Mito translation |  |  |  |  |  |
| F55C5.5 | tsfm-1 | Mitochondrial translation elongation factor EF-Tsmt | Metabolism: Mito translation |  |  |  |  |  |
| W10D9.5 |  | Translocase of outer mitochondrial membrane complex, subunit TOM22 | Metabolism: Mitochondial |  |  |  |  |  |
| T09B4.9 |  | Mitochondrial import inner membrane translocase, subunit TIM44 | Metabolism: Mitochondial |  |  |  |  |  |
| F23H12.2 |  | Translocase of outer mitochondrial membrane complex, subunit TOM20 | Metabolism: Mitochondial |  |  |  |  |  |
| F01G4.6 |  | Mitochondrial phosphate carrier protein | Metabolism: Mitochondial |  |  |  |  |  |
| F54B3.3 | atad-3 | AAA+-type ATPase; mito inner membrane | Metabolism: Mitochondial |  |  |  |  |  |
| T20G5.2 | cts-1 | a citrate synthase, predicted to be mitochondrial | Metabolism |  |  |  |  |  |
| H14A12.2 | fum-1 | fumarase (mitochondrial) | Metabolism |  |  |  |  |  |
| M01F1.3 |  | lipoate synthase | Metabolism |  |  |  |  |  |
| T02G5.7 |  | Acetyl-CoA acetyltransferase | Metabolism |  |  |  |  |  |
| C49F5.1 | sams-1 | S-adenosyl methionine synthetase | Metabolism | [Extended lifespan](http://wormbase.org/db/misc/paper?name=WBPaper00026715;class=Paper) |  |  |  |  |
| [F20C5.1](http://www.wormbase.org/db/gene/gene?name=WBGene00004051;class=Gene) | pme-3 | Poly(ADP-ribose) glycohydrolase | Metabolism |  |  |  |  |  |
| [AC8.1](http://www.wormbase.org/db/seq/sequence?name=AC8.1;class=Gene_name) | [pme-6](http://www.wormbase.org/db/gene/gene?name=WBGene00004054;class=Gene) | poly(ADP-ribose) metabolism enzyme | Metabolism |  |  |  |  |  |
| LLC1.3 |  | dihydrolipoamide dehydrogenase, predicted to be mitochondrial | Metabolism |  |  |  |  | paraquat resistance; |
| F54H12.1 | aco-2 | aconitase, predicted to be mitochondrial | Metabolism |  | lifespan extended |  |  |  |
| [C56G2.6](http://www.wormbase.org/db/seq/sequence?name=C56G2.6;class=Gene_name) | let-767 | steroid dehydrogenase | Metabolism |  |  |  | unc-54:POLYQ(Q35)YFP aggregation |  |
| T06D8.6 | cchl-1 | Holocytochrome c synthase/heme-lyase | Metabolism | [Extended lifespan](http://wormbase.org/db/misc/paper?name=WBPaper00026715;class=Paper) |  |  |  |  |
| Y39B6A.3 |  | Fe-S cluster biosynthesis protein ISA1, predicted to be mitochondrial | Metabolism |  |  |  |  |  |
| Y49E10.2 | glrx-5 | Glutaredoxin-related protein, predicted to be mitochondrial | Metabolism |  |  |  |  |  |
| Y47G6A.10 | spg-7 | nuclear-encoded mitochondrial metalloprotease, predicted to regulate degradation of mito proteins | Metabolism |  |  | [extended lifespan; POLYQ(Q40)YPF loc; sod-3:GFP; DAF-16:GFP](http://wormbase.org/db/misc/paper?name=WBPaper00029254;class=Paper) | unc-54:POLYQ(Q35)YFP aggregation |  |
| T24H7.1 | phb-2 | one of two subunits of the mitochondrial prohibitin complex | Metabolism |  |  |  |  |  |
| K01C8.7 |  | Mitochondrial FAD carrier protein | Metabolism |  |  |  |  |  |
| D2013.5 | eat-3 | mitochondrial dynamin-related protein | Metabolism |  |  |  |  |  |
| T20F5.2 | pbs-4 | B-type subunit of the 26S proteasome core particle | Protein folding and turnover |  |  |  | unc-54:POLYQ(Q35)YFP aggregation |  |
| R12E2.3 | rpn-8 | RPN8 subunit of 26S proteasome | Protein folding and turnover |  |  |  | unc-54:POLYQ(Q35)YFP aggregation |  |
| Y110A7A.14 | pas-3 | PSMA4 subunit of 20S proteasome | Protein folding and turnover |  |  |  |  |  |
| F56H1.4 | rpt-5 | 26S proteasome regulatory complex, ATPase RPT5 | Protein folding and turnover |  |  |  | unc-54:POLYQ(Q35)YFP aggregation |  |
| C36B1.4 | pas-4 | 20S proteasome, regulatory subunit PSMA7 | Protein folding and turnover |  |  |  | unc-54:POLYQ(Q35)YFP aggregation |  |
| C47B2.4 | pbs-2 | 20S proteasome, regulatory subunit beta type PSMB7 | Protein folding and turnover |  |  |  | unc-54:POLYQ(Q35)YFP aggregation |  |
| K05C4.1 | pbs-5 | 20S proteasome, regulatory subunit beta type PSMB5 | Protein folding and turnover |  |  |  | unc-54:POLYQ(Q35)YFP aggregation |  |
| ZK20.5 | rpn-12 | non-ATPase subunit of 19S proteasome complex | Protein folding and turnover |  |  |  |  |  |
| T06D8.8 | rpn-9 | non-ATPase subunit of 19S proteasome complex | Protein folding and turnover |  |  |  |  |  |
| K07D4.3 | rpn-11 | non-ATPase subunit of 19S proteasome complex | Protein folding and turnover |  |  |  | unc-54:POLYQ(Q35)YFP aggregation |  |
| F23F1.8 | rpt-4 | ATPase subunit of 19S proteasome complex | Protein folding and turnover |  |  |  |  |  |
| C23G10.4 | rpn-2 | ATPase subunit of 19S proteasome complex | Protein folding and turnover |  |  |  |  |  |
| F23F12.6 | rpt-3 | ATPase subunit of 19S proteasome complex | Protein folding and turnover |  |  |  |  |  |
| F57B9.10 | rpn-6 | non-ATPase subunit of 19S proteasome complex | Protein folding and turnover |  |  |  | unc-54:POLYQ(Q35)YFP aggregation |  |
| C02F5.9 | pbs-6 | 20S proteasome, regulatory subunit beta type PSMB1 | Protein folding and turnover |  |  |  |  |  |
| C30C11.2 | rpn-3 | 26S proteasome regulatory complex, subunit RPN3 | Protein folding and turnover |  |  |  |  |  |
| Y49E10.1 | rpt-6 | 26S proteasome regulatory complex, ATPase RPT6 | Protein folding and turnover |  |  |  |  |  |
| [F49C12.8](http://www.wormbase.org/db/seq/sequence?name=F49C12.8;class=Gene_name) | rpn-7 | 26S proteasome regulatory complex, subunit RPN7 | Protein folding and turnover |  |  |  |  |  |
| T22D1.9 | rpn-1 | 26S proteasome regulatory complex, subunit RPN1 | Protein folding and turnover |  |  |  |  |  |
| K08D12.1 | pbs-1 | 20S proteasome, regulatory subunit beta type PSMB6 | Protein folding and turnover |  |  |  | unc-54:POLYQ(Q35)YFP aggregation |  |
| C52E4.4 | rpt-1 | 26S proteasome regulatory complex, ATPase RPT1 | Protein folding and turnover |  |  |  |  |  |
| [CD4.6](http://www.wormbase.org/db/seq/sequence?name=CD4.3;class=Gene_name) | pas-6 | 20S proteasome, regulatory subunit beta type PSMA1 | Protein folding and turnover |  |  |  | unc-54:POLYQ(Q35)YFP aggregation |  |
| F39H11.5 | pbs-7 | 20S proteasome, regulatory subunit beta type PSMB4 | Protein folding and turnover |  |  |  |  |  |
| [Y38F2AR.8](http://www.wormbase.org/db/seq/sequence?name=Y38F2AR.8;class=Gene_name) | ppgn-1 | paraplegin homolog | Protein folding and turnover |  |  |  |  |  |
| F08G12.4 | vhl-1 | von Hippel-Lindau tumor suppressor, E3 ligase | Protein folding and turnover |  |  |  |  |  |
| Y82E9BR.3&Y82E9BR.15 |  | 2 genes: (1) Mitochondrial ATP synthase (2) elongin C | Protein folding and turnover |  |  |  |  |  |
| H38K22.2 | dcn-1 | required for neddylation of CUL-3 cullin | Protein folding and turnover |  |  |  |  |  |
| F45H11.2 | ned-8 | Ubiquitin-like protein | Protein folding and turnover |  |  |  |  |  |
| ZK1240.1 |  | Predicted E3 ubiquitin ligase | Protein folding and turnover |  |  |  |  |  |
| C18F10.7 |  | Ankyrin repeat protein with ubiquitin interaction motif | Protein folding and turnover |  |  |  |  |  |
| E03H4.8 |  | Vesicle coat complex COPI, beta subunit | Vesicular transport |  |  |  |  |  |
| Y105E8A.9 | apg-1 | Vesicle coat complex AP-1, gamma subunit | Vesicular transport |  |  |  | unc-54:POLYQ(Q35)YFP aggregation |  |
| F09E5.11 |  | divergent ortholog of S. cerevisiae VPH2 | Vesicular transport |  |  |  |  |  |
| C30A5.5 | snb-5 | synaptobrevin-like protein | Vesicular transport |  |  |  |  |  |
| K02D10.5 | snap-29 | SNAP component of SNARE complex | Vesicular transport |  |  |  | unc-54:POLYQ(Q35)YFP aggregation |  |
| T20G5.1 | chc-1 | Vesicle coat protein clathrin, heavy chain | Vesicular transport |  |  |  |  |  |
| F38E11.5 |  | Vesicle coat complex COPI, beta subunit | Vesicular transport |  |  |  |  |  |
| [C17H12.14](http://www.wormbase.org/db/gene/gene?name=WBGene00006917;class=Gene) | [vha-8](http://www.wormbase.org/db/gene/gene?name=WBGene00006917;class=Gene) | Vacuolar-type H^+^-ATPase component | Transporters and Channels |  |  |  | unc-54:POLYQ(Q35)YFP aggregation |  |
| Y55H10A.1 | vha-19 | Vacuolar-type H^+^-ATPase accessory subunit | Transporters and Channels |  |  |  |  |  |
| [Y38F2AL.4](http://www.wormbase.org/db/seq/sequence?name=Y38F2AL.4;class=Gene_name) | vha-3 | Vacuolar-type H^+^-ATPase component | Transporters and Channels |  |  |  |  |  |
| [Y38F2AL.3](http://www.wormbase.org/db/gene/gene?name=WBGene00006920;class=Gene) | vha-11 | Vacuolar-type H^+^-ATPase component | Transporters and Channels |  |  |  |  |  |
| Y38F2AR.2 | trap-3 | Translocon-associated complex gamma subunit | Transporters and Channels |  |  |  |  |  |
| C14C10.4 |  | Uncharacterized | Transporters and Channels |  |  |  |  |  |
| B0285.6 |  | Sodium-coupled carboxylate transporter | Transporters and Channels |  |  |  |  |  |
| F23F12.3 |  | Major facilitator superfamily of tranporters | Transporters and Channels |  |  |  |  |  |
| Y82E9BR.16 |  | Major facilitator superfamily of tranporters | Transporters and Channels |  |  |  |  |  |
| F29G9.3 | aps-1 | Clathrin adaptor complex, small subunit | Transporters and Channels |  |  |  |  |  |
| F14F4.3 | mrp-5 | CFTR homolog / ABC superfamily of transporters | Transporters and Channels |  |  |  |  |  |
| W07A12.6 | oac-54 | transmembrane acyltransferase, homologous to rhy-1 | Signal transduction and cytoskeleton |  |  |  |  |  |
| W07A12.7 | rhy-1 | transmembrane acyltransferase; regulator of HIF-1-mediated txn | Signal transduction and cytoskeleton |  |  |  |  |  |
| F47B3.1 |  |  | Signal transduction and cytoskeleton |  |  |  |  |  |
| F23H11.5 |  | Uncharacterized transmembrane protein | Signal transduction and cytoskeleton |  |  |  |  |  |
| F58B6.2 | inft-1 | formin-related protein | Signal transduction and cytoskeleton |  |  |  |  |  |
| R10E4.9 |  | Protein tyrosine phosphatase-like protein | Signal transduction and cytoskeleton |  |  |  |  |  |
| B0222.5 |  | Serine proteinase inhibitor (KU family) | Signal transduction and cytoskeleton |  |  |  |  |  |
| C14B9.2 |  | Protein disulfide isomerase | Signal transduction and cytoskeleton |  |  |  |  |  |
| F22E12.4 | egl-9 | dioxygenase that negatively regulates HIF-1 | Signal transduction and cytoskeleton |  |  |  |  |  |
| T12A2.13 | srg-6 | seven transmembrane-domain receptor | Signal transduction and cytoskeleton |  |  |  |  |  |
| T19B4.4 | dnj-21 | molecular chaperone (DnaJ superfamily) | Protein folding and turnover |  |  |  |  |  |
| C14B9.1 | hsp12.2 | small heat-shock protein | Protein folding and turnover |  |  |  |  |  |
| Y54E10B_154.c | dnj-28 | DnaJ superfamily | Protein folding and turnover |  |  |  |  |  |
| T19E7.2 | [skn-1](http://www.wormbase.org/db/gene/gene?name=WBGene00020961;class=Gene) | Transcription factor homologous to mammalian NRF | Transcription/translation/DNA/RNA processing |  |  |  |  |  |
| [Y55F3AM.12](http://www.wormbase.org/db/gene/gene?name=WBGene00021929;class=Gene) | dcap-1 | homologous to mRNA decapping components | Transcription/translation/DNA/RNA processing |  |  |  |  |  |
| C46G7.1 |  | ribonuclease, transmembrane domain | Transcription/translation/DNA/RNA processing |  |  |  |  |  |
| R144.2 | pcf-11 | mRNA cleavage and polyadenylation factor I/II complex, subunit Pcf11 | Transcription/translation/DNA/RNA processing |  |  |  |  |  |
| C29H12.1 | rrt-2 | Arginyl-tRNA synthetase | Transcription/translation/DNA/RNA processing |  |  |  |  |  |
| Y57E12AL.5 | mdt-6 | transcriptional mediator of the Med6 family | Transcription/translation/DNA/RNA processing |  |  |  |  |  |
| F38A1.8 |  | Signal recognition particle receptor, alpha subunit | Transcription/translation/DNA/RNA processing |  |  |  |  |  |
| Y47D3B.7 | sbp-1 | homologous to SREBP transcription factor | Transcription/translation/DNA/RNA processing |  |  |  | unc-54:POLYQ(Q35)YFP aggregation |  |
| Y47D3A.26 | smc-3 | cohesin subunit; roles in chromosome segregation | Transcription/translation/DNA/RNA processing |  |  |  |  |  |
| ZK1058.2 | pat-3 | integrin beta subunit | Signal transduction and cytoskeleton |  |  |  |  |  |
| K07C5.1 | arx-2 | Actin-related protein Arp2 | Signal transduction and cytoskeleton |  |  |  |  |  |
| W07B3.2 | gei-4 | coiled-coil domain, regulation of intermediate filament dynamics | Signal transduction and cytoskeleton |  |  |  |  |  |
| Y54G11A.8 | ddl-3 | tetratricolpeptide-repeat protein interaction motif | Signal transduction and cytoskeleton | lifespan extended |  |  |  |  |
| C46H11.6 |  | PDZ domain containing protein | Other |  |  |  |  |  |
| C29H12.2 |  | integral membrane protein, homology to transporters | Other |  |  |  |  |  |
| K04G7.11 |  | homology to pre-mRNA splicing factor SYF2 | Other |  |  |  |  |  |
| F49C12.11 |  | uncharacterized coiled-coil domain protein | Other |  |  |  |  |  |
| ZK856.12 |  | uncharacterized protein | Other |  |  |  |  |  |
| R155.4 |  | WSN domain (worm-specific N-terminal domain) | Other |  |  |  |  |  |
| F26F4.9 |  | uncharacterized protein | Other |  |  |  |  |  |
| C14C10.3 | ril-2 | uncharacterized protein | Other | lifespan extended |  |  |  |  |
| C53A5.1 | ril-1 | uncharacterized protein | Other | lifespan extended |  |  | unc-54:POLYQ(Q35)YFP aggregation |  |
| K12H4.5 |  | uncharacterized protein | Other |  |  |  |  | paraquat resistance; |
| [Y38E10A.24](http://www.wormbase.org/db/seq/sequence?name=Y38E10A.24;class=Gene_name) |  | uncharacterized protein | Other |  |  |  |  |  |
| B0491.5 |  | uncharacterized protein | Other |  |  |  |  |  |
| Y54G9A.5 |  | uncharacterized intregal membrane protein | Other |  |  |  |  |  |
| F29C4.2 |  | similarity to cytochrome c oxidase subunit Vic | Other |  |  |  |  | paraquat resistance; |
| [F38A5.5](http://www.wormbase.org/db/gene/gene?name=WBGene00018162;class=Gene) | nspb-3 | nematode-specific peptide family | Other |  |  |  |  |  |
| F44E5.1 |  | uncharacterized protein | Other |  |  |  |  |  |
