## Supplement Figure 2 for "Crosstalk in oxygen homeostasis networks: SKN-1/NRF inhibits the HIF-1 hypoxia-inducible factor in *Caenorhabditis elegans*"

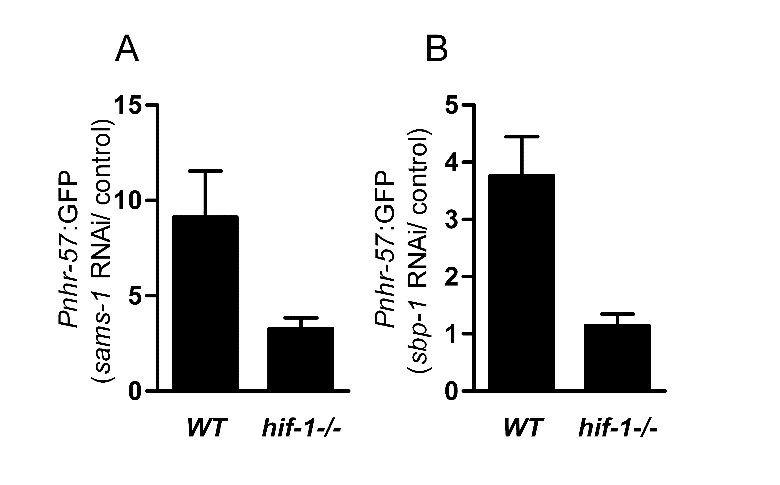

**S2 Fig. RNAi inactivation of *sams-1* or *sbp-1* increased *Pnhr-57:GFP* expression.**

(A) RNAi for *sams-1* (S-adenosyl methionine synthetase) increased *Pnhr‑57*:*GFP* expression more than 7-fold in animals carrying the wild-type *hif-1* allele relative to control RNAi, and increased the reporter 3-fold in animals carrying the *hif-1(ia04)* deletion. The difference in RNAi effect between *hif-1(+)* and *hif-1(ia04)* strains is statistically significant (**p* < 0.05, from six independent experiments, by student *t-*test). (B) RNAi for the SREBP homolog *sbp-1* increased expression of the reporter more than 3-fold in animals carrying the wild-type *hif-1* allele, but had no effect on *Pnhr-57*:*GFP* expression in *hif-1(ia04)* mutants. The difference in RNAi effect between *hif-1(+)* and *hif-1(ia04)* strains is statistically significant (**p* < 0.05, from five independent experiments, by student *t-*test). GFP levels were determined by protein blots, and the control animals were fed on bacteria carrying the empty RNAi vector (L4440). The experiments were conducted in RNAi-sensitive strains (*rrf-3(pk1426*)).
