## Supplement Table 2 for "Crosstalk in oxygen homeostasis networks: SKN-1/NRF inhibits the HIF-1 hypoxia-inducible factor in *Caenorhabditis elegans*"

**S2 Table. Genes for which RNAi caused *hif-1*-dependent increase of *Pnhr-57:GFP* expression.**

|  | **Gene** | **Description** |
| --- | --- | --- |
| Mitochondrial/ Metabolism | T09B4.9 | Mitochondrial import inner membrane translocase |
|  | *sdhb-1* | Succinate dehydrogenase subunit |
|  | *sams-1* | S-adenosyl methionine synthetase |
|  | *sco-1* | Putative cytochrome C oxidase assembly protein |
|  | W02F12.5 | oxoglutarate dehydrogenase complex |
| Protein turnover | *rpn-11* | Proteasome regulatory particle |
|  | *rpn-12* | Proteasome regulatory particle |
| Transcription/ translation | *rrt-2* | Arginyl-tRNA synthetase |
|  | *sbp-1* | Transcription factor: SREBP homolog |
|  | *skn-1* | Transcription factor: Nrf homolog |
| Known regulators of HI | *egl-9* | Prolyl hydroxylase |
|  | *rhy-1* | Acyltransferase |
|  | *vhl-1* | E3 ligase, Von Hipple Lindau |
